# To what extent can life history strategies inform species conservation planning?

**DOI:** 10.1101/2024.12.09.626826

**Authors:** Emily A. Stevenson, Sol Lucas, Philip J. K. McGowan, Isabel M. Smallegange, Louise Mair

**Author notes:** Corresponding author and mailing address: Emily Stevenson, Modelling, Evidence and Policy Group, School of Natural and Environmental Sciences, Agriculture Building, King’s Road, Newcastle University, Newcastle upon Tyne, NE1 4LB, UK. I.M. Smallegange and L. Mair should be considered joint senior author.

## Abstract

Global policy aims to prevent species extinctions; to support these aims conservation planners must effectively target interventions to reduce the extinction risk of species. However, there is often a lack of knowledge on the magnitude and direction of species responses to interventions and in turn the extent to which a species extinction risk is reduced. If we can use a species’ life history strategies to predict their responses to interventions, this offers a promising approach to better understand species extinction risks and conservation potential. Here we apply Dynamic Energy Budget Integral Project Models to 23 reptile species to investigate whether their life history traits can be summarised into a life history strategy framework using principal component analysis, and whether species positions along these axes predict their population growth rate, demographic resilience, sensitivity to perturbations and extinction risk. We found that species positions on reproductive and pace of life axes predicted reptile population growth rate and demographic resilience but not sensitivity to perturbations or extinction risk. Our findings show that reptile life history strategies can inform our understanding of reptile species conservation potential and could be applied to influence management decisions such as establishing monitoring timelines.

## 1. Introduction

In 2022, the Kunming-Montreal Global Biodiversity Framework (KMGBF) was agreed by Parties to the Convention on Biological Diversity (CBD, 2022). The KMGBF contains global goals to halt and reverse biodiversity loss by 2050, including Goal A which stipulates that, for species specifically, “Human induced extinction of known threatened species is halted, and, by 2050, the extinction rate and risk of all species are reduced tenfold” (CBD, 2022). Of the 166,061 species currently documented on the International Union for Conservation of Nature Red List of Threatened Species (hereafter IUCN Red List), more than 46,000 species are classified as threatened with extinction (IUCN, 2024), which has been extrapolated to an estimated one million species at risk of extinction globally (IPBES, 2019; <u>Purvis & Isbell</u> <u>2022)</u>. Moreover, during the 10-years preceding the agreement of the KMGBF, at least four vertebrate species went extinct (IUCN, 2024).

There is evidence that conservation can prevent species extinctions; expert elicitation estimated that at least 28-48 birds and mammal species were prevented from going extinct because of conservation actions between 1993-2020 (Bolam et al. 2021), and at least 148 species of ungulates would have deteriorated by at least one IUCN Red List extinction risk category without conservation actions (Hoffmann et al. 2015). Yet, with thousands of species listed as requiring targeted active management to avert human induced extinction (Bolam et al. 2023), achieving the total prevention of human-induced extinction and the overall reduction of extinction risk will require effective planning, implementation and upscaling of conservation actions (Crees et al. 2016; Nicol et al. 2019).

To ensure the effective and efficient allocation of resources to prevent extinctions, it is necessary to understand why some actions succeed while others fail (Crees et al. 2016; Catalano et al. 2019). One of the major challenges is a lack of knowledge on the magnitude and direction of species responses to interventions, which can hinder planning and implementation, and restrict the effectiveness of conservation interventions (Grainger et al. 2019; Nicol et al. 2019; Christie et al. 2020). Robust monitoring of species’ status and response to interventions is thus essential but can be costly and time-consuming (McDonald-Madden et al. 2010; Moussy et al. 2022; Stephenson et al. 2022) and species’ populations can take a long time to respond to interventions (Watts et al. 2020; Piipponen-Doyle et al. 2021). In some scenarios, waiting for monitoring to gather all desirable information may not be an option when conservation intervention is urgently needed (Grainger et al. 2019). There is thus a need for alternative, efficient approaches that can inform conservation planning based on our existing species knowledge.

One approach to inform conservation planning is to use life history data. The most commonly observed life history strategy of species is the fast-slow continuum, where life history strategies are structured along a single axis defined by the trade-off between survival and reproduction, where “fast” species mature early and have high reproduction but low survival and “slow” species mature later in life and have low reproduction but high survival (Gaillard et al. 2016). The position of a species on this axis has been linked to conservation status, as slow-lived bird and mammal species have higher extinction rates (Cooke et al. 2019), and slow-lived mammals display more negative responses to disturbances (Suraci et al. 2021). A second major life history axis is the reproductive strategy axis (Salguero-Gómez et al. 2016), which is characterised by high lifetime reproductive output and high mortality at one end, and low reproductive output and low mortality at the other (in some cases this second axis relates to development instead; Stott *et al*., 2024). At one extreme end of this axis are semelparous species (single reproductive event in their lifespan) with low variability in mortality, and then iteroparous species (multiple reproductive cycles throughout their lifespan) with high variability in mortality are ranked at the other extreme end (Healy et al. 2019).

Together, these two axes comprise the fast-slow continuum and reproductive strategy framework (Salguero-Gómez et al. 2016), within which a species’ position along these axes can predict its population performance, demographic resilience, and even extinction risk (Salguero-Gómez et al. 2016; Salguero-Gómez 2017). Furthermore, by conducting a perturbation analysis, where the influence of a proportional change in each life history parameter is compared to the resulting change in a species population growth rate, it is possible to identify the most sensitive life history stage of the species and how that changes with the environment (Smallegange & Lucas 2024). A species’ position within the fast-slow continuum and reproductive strategy framework could also predict this sensitivity to environmental change (Smallegange & Lucas 2024). Using these findings, it may be possible to reduce uncertainty about how species may respond to conservation interventions based on their life history strategy, and identify interventions that are more or less likely to succeed based on the life history stage being targeted (Salguero-Gómez 2017; Warret Rodrigues et al. 2021).

One species group that could benefit from the additional insights gained from life history strategy analyses are reptiles, which are underrepresented among vertebrate species in conservation literature (Roll et al. 2017; Di Marco et al. 2017). The IUCN Red List recently completed a comprehensive assessment of reptiles and found that 18% (1,845 out of 10,311 species) are threatened with extinction (Cox *et al*., 2022; IUCN 2024). Life-history strategies have been studied in individual reptiles species in relation to conservation (Briggs-Gonzalez et al. 2017), but the fast-slow continuum and reproductive strategy framework has not been applied across the Reptilia class. By applying this approach, it may be possible to help improve the development of targeted conservation interventions for reptiles to reduce extinction risk (Roll et al. 2017).

Here we investigate to what extent life history strategies could contribute to our understanding of species conservation potential, and how this understanding could be used to improve conservation planning. We explore if and how reptile demographic rates can be structured into a life history strategy framework, and whether reptile species’ position along these axes predict their population performance, demographic resilience, sensitivity to perturbations and risk of extinction. To this end, we parameterise Dynamic Energy Budget Integral Projection Models (DEB-IPMs), which include an energy budget model that describes a species’ growth and reproduction, where energy intake is determined by setting an experienced feeding level (Smallegange et al. 2017). DEB-IPMs are ideal for modelling demographic data in the context of this study because they can describe the demography of species using only eight parameters (Smallegange et al. 2017). This limited data requirement is beneficial for conservation planners, allowing them to make informed decisions on interventions when working with data-sparse species.

To achieve our aim, we (objective 1) parameterised DEB-IPMs for 23 reptile species taken from the DEBBIES dataset (Smallegange & Lucas 2024) to calculate a set of representative life history traits based on schedules of survival, growth, and reproduction (Smallegange et al. 2017). We then evaluated the variation in these traits along major axes using a mass-corrected principal component analysis (PCA). Next, we (objective 2) tested whether the position of species along these axes predicted variables of conservation interest: population performance, demographic resilience, sensitivity to perturbations and risk of extinction. Our results allowed us to identify significant associations between life history strategies and variables of conservation interest, suggest possible applications of derived life history traits and propose future applications of life history data that may benefit conservation planning.

## 2. Methods

A DEB-IPM is a population model that tracks cohorts of female individuals in a population, capturing the demographic rates of survival, growth, reproduction and the relationship between parent and offspring size, using eight life history parameters (Figure 1; Smallegange et al. 2017; Smallegange & Lucas 2024). This model also includes the influence of environmentally driven variation in feeding levels in the trade-off individuals make when allocating energy for growth or reproduction (Smallegange et al. 2017; Smallegange & Lucas 2024). A brief description of the model can be found in the appendix, and full details of how to parameterise and apply DEB-IPMs are in Smallegange and Lucas (2024).

**Figure 1:**
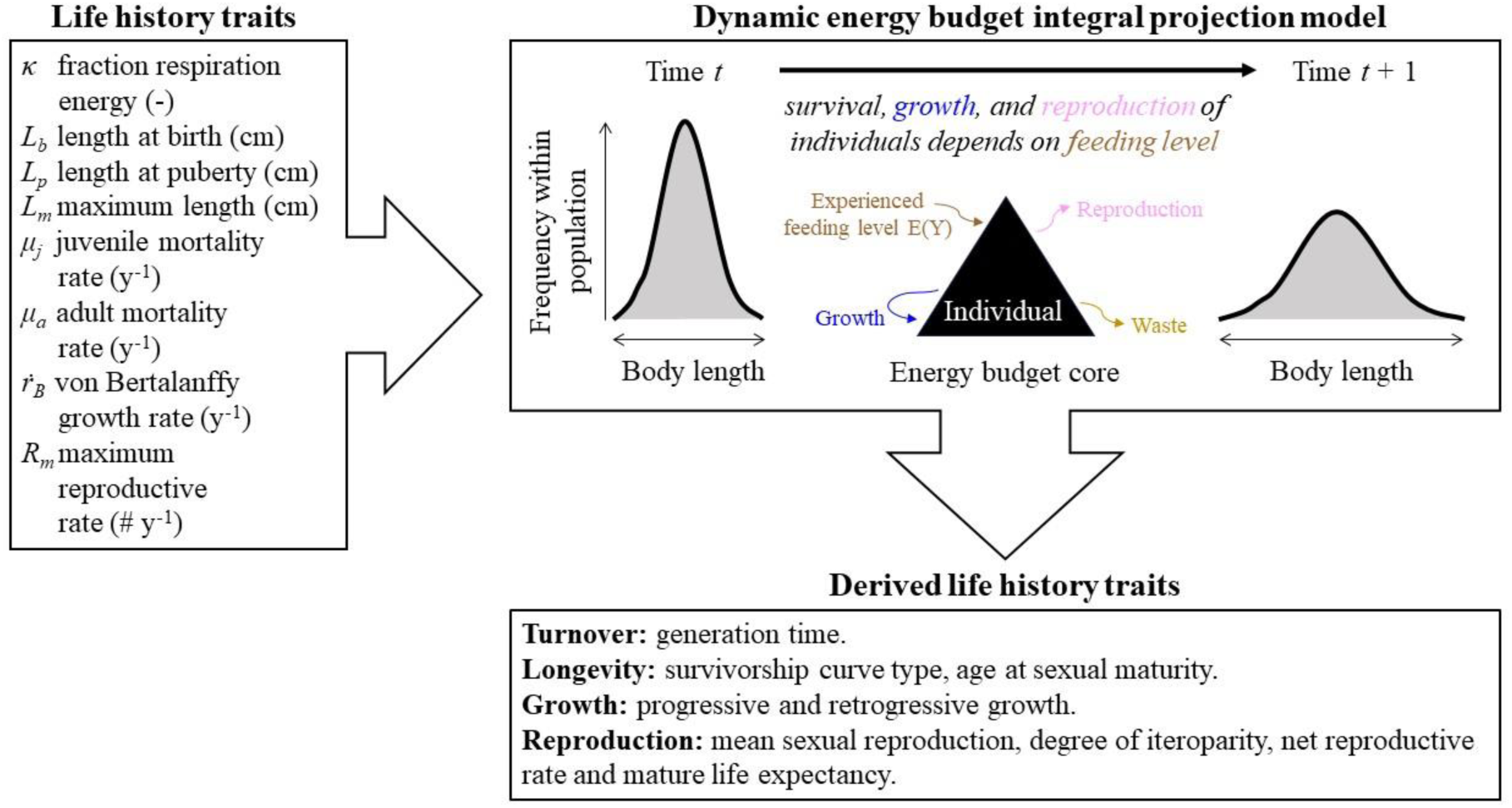
Workflow of parameterising a DEB-IPM using the DEBBIES database (Smallegange & Lucas 2024). Eight life history trait values are required to parameterise a DEB-IPM (top-left box). Once parameterised, it can be used to calculate a further nine derived life history traits (Table 2) that in turn can be summarised into life history strategies (Smallegange & Lucas 2024).

### 2.1. Identifying life history strategies (objective 1)

The DEBBIES dataset (Version 5) contains estimates of the eight DEB-IPM life history parameters for many ectothermic species (Smallegange & Lucas 2024). We extracted these records for all 23 reptile species listed in DEBBIES (Table 1), which allowed for a representative selection of reptile species, including all extant orders, and a mixture of threatened and non-threatened species. Among the 184 individual parameter estimates obtained (eight estimates for each of the 23 species), we updated 19 using published records from primary literature (Table S1). We also gathered data on the extinction risk of each species using the IUCN Red List (IUCN, 2024), and maximum female body mass by searching primary literature.

**Table 1:** DEB-IPM model parameters for females of each species, taken from the DEBBIES Dataset (Smallegange & Lucas 2024). Extinction risk data were gathered from the IUCN Red List (2024). The parameter *κ* (-) indicates the fraction of energy allocated to respiration as opposed to reproduction, *L_b_* indicates length at birth (cm), *L_p_* indicates length at puberty (cm), *L_m_* indicates maximum length (cm), *μ_j_* indicates juvenile mortality rate (y^-1^), *μ_a_* indicates adult mortality rate (y^-1^), *ṙ_B_* indicates von Bertalanffy growth rate (y^-1^), and *R_m_* indicates maximum reproductive rate (# y^-1^) (See also Fig. 1). Mass values are in kg. All changes and references are shown in Table S1.

| Species | Common name | Extinction Risk Category | $\kappa$ (-) | $L_b$ (cm) | $L_p$ (cm) | $L_m$ (cm) | $\mu_j$ ($y^{-1}$) | $\mu_a$ ($y^{-1}$) | $\dot{r}_B$ ($y^{-1}$) | $R_m$ ($\# y^{-1}$) | Mass (kg) |
| --- | --- | --- | --- | --- | --- | --- | --- | --- | --- | --- | --- |
| *Caiman crocodilus* | Spectacled caiman | Least concern | 0.95 | 13.00 | 120.00 | 161.00 | 0.16 | 0.16 | 0.13 | 40.00 | 20.000 |
| *Crocodylus johnstoni* | Australian freshwater crocodile | Least concern | 0.97 | 24.00 | 131.00 | 200.00 | 0.78 | 0.04 | 0.10 | 18.00 | 30.000 |
| *Crocodylus niloticus* | Nile crocodile | Least concern | 0.66 | 29.00 | 280.00 | 427.00 | 0.12 | 0.12 | 0.07 | 95.01 | 94.200 |
| *Gavialis gangeticus* | Gharial | Critically endangered | 0.74 | 37.00 | 300.00 | 375.00 | 0.12 | 0.12 | 0.09 | 37.01 | 150.000 |
| *Sphenodon punctatus* | Tuatara | Least concern | 0.79 | 10.50 | 34.80 | 80.00 | 0.27 | 0.05 | 0.05 | 6.00 | 0.690 |
| *Boa constrictor* | Red-tailed boa | Least concern | 0.62 | 45.00 | 170.00 | 450.00 | 0.12 | 0.12 | 0.05 | 65.01 | 27.000 |
| *Egernia striolata* | Tree-crevice Skink | Least concern | 0.65 | 5.17 | 10.00 | 12.10 | 0.40 | 0.40 | 0.75 | 4.00 | 0.030 |
| *Eunectes murinus* | Green anaconda | Least concern | 0.34 | 37.00 | 302.00 | 914.00 | 0.12 | 0.12 | 0.11 | 100.01 | 74.000 |
| *Natrix helvetica* | European grass snake | Least concern | 0.59 | 17.00 | 65.00 | 130.00 | 0.42 | 0.40 | 0.11 | 62.00 | 0.300 |
| *Takydromus*<br>*hsuehshanensis* | Snow Mountain<br>grasslizard | Least concern | 0.66 | 2.60 | 5.40 | 7.20 | 0.40 | 0.40 | 0.48 | 6.00 | 0.047 |
| *Tiliqua rugosa* | Shingleback lizard | Least concern | 0.91 | 16.50 | 23.00 | 32.80 | 1.06 | 0.11 | 0.50 | 1.70 | 0.700 |
| *Chelodina*<br>*oblonga* | Northern snake-necked turtle | Near threatened | 0.82 | 3.10 | 21.00 | 30.00 | 0.02 | 0.02 | 0.22 | 14.00 | 2.285 |
| *Cuora*<br>*flavomarginata* | Yellow-margined box<br>turtle | Endangered | 0.58 | 4.20 | 14.60 | 16.70 | 0.15 | 0.15 | 0.10 | 9.00 | 0.976 |
| *Podocnemis*<br>*unifilis* | Yellow-spotted river<br>turtle | Vulnerable | 0.85 | 4.80 | 31.00 | 52.00 | 0.08 | 0.08 | 0.18 | 40.00 | 6.900 |
| *Dermochelys*<br>*coriacea* | Leatherback turtle | Vulnerable | 0.92 | 7.00 | 121.00 | 175.00 | 0.46 | 0.11 | 0.12 | 200.53 | 650.000 |
| *Chelonia mydas* | Green turtle | Endangered | 0.72 | 5.00 | 80.00 | 99.00 | 0.13 | 0.05 | 0.09 | 110.00 | 243.500 |
| *Caretta caretta* | Loggerhead turtle | Vulnerable | 0.65 | 4.50 | 80.00 | 125.00 | 0.24 | 0.09 | 0.12 | 280.00 | 107.000 |
| *Eretmochelys*<br>*imbricata* | Hawksbill turtle | Critically<br>endangered | 0.80 | 4.13 | 78.60 | 105.00 | 0.24 | 0.06 | 0.09 | 288.00 | 112.200 |
| *Chelydra*<br>*serpentina* | Snapping turtle | Least concern | 0.80 | 2.70 | 22.00 | 39.70 | 0.46 | 0.21 | 0.11 | 69.00 | 12.800 |
| *Kinosternon*<br>*flavescens* | Yellow mud turtle | Least concern | 0.8 | 2.19 | 8.83 | 11.70 | 0.54 | 0.06 | 0.11 | 9.00 | 0.349 |
| *Gehyra*<br>*variegata* | Varied dtella | Least concern | 0.8 | 2.80 | 4.80 | 5.90 | 1.61 | 0.93 | 0.85 | 2.00 | 0.040 |
| *Sceloporus*<br>*undulatus* | Eastern fence lizard | Least concern | 0.8 | 2.50 | 4.70 | 8.00 | 2.81 | 1.18 | 0.41 | 40.00 | 0.018 |
| *Thamnophis*<br>*elegans* | Terrestrial gartersnake | Least concern | 0.80 | 18.77 | 40.00 | 70.00 | 1.11 | 0.29 | 0.40 | 14.35 | 0.117 |

Once parametrised, we calculated nine life history traits (Table 2) in MatLab (using publicly available code from Smallegange and Lucas (2024)). We tested a range of feeding levels (*E(Y)*) and found that four species did not demonstrate a positive population growth rate at any feeding level, meaning they always went extinct regardless of environmental quality. As we were interested in exploring species growth and recovery these species were therefore excluded from the analysis, leaving 19 species. We selected a feeding level of *E*(*Y*)= 0.9 for all further analyses, because this was the lowest feeding level at which all remaining 19 species survived. This was preferred to the alternative of setting the feeding level at *E*(*Y*)= 1.0, as this represents an individual with a full gut, which is not as realistic given the natural variability in a species’ environment (Smallegange et al. 2020).

**Table 2:** Derived life history traits that can be generated using DEB-IPM.

| Life History Trait | Symbol | Definition | Formula |
| --- | --- | --- | --- |
| Generation time | $T$ | Number of years required for the individuals of a population to be fully replaced by new ones. | $T = \frac{\log(R_0)}{\log(\lambda)}$ |
| Survivorship curve | $H$ | Keyfitz' entropy ($H < 1$ denotes increasing mortality rate with age, $H = 1$ denotes a constant mortality rate with age, and $H > 1$ denotes decreasing mortality rate with age). | $H = - \frac{\sum_{x=0}^{x=\eta_e} \log(l_x) l_x}{\sum_{x=0}^{x=\eta_e} l_x}$ |
| Age at maturity | $L_a$ | Number of years that it takes an average individual in the population to become reproductive. | $L_a = \eta_{L_b}$ |
| Progressive growth | $\gamma$ | Mean probability of growing to a larger length across the length domain $\Omega$. | $\gamma = \sum_i^m \bar{G}_{i,j} \vert _{i < j}$ |
| Retrogressive growth | $\rho$ | Mean probability of growing to a smaller length across the length domain $\Omega$. | $\rho = \sum_i^m \bar{G}_{i,j} \vert _{i > j}$ |
| Mean recruitment success | $\varphi$ | Mean per-capita number of recruits across the length domain $\Omega$. | $\varphi = \sum_i^m \bar{V}_{i,j}$ |
| Degree of iteroparity | $S$ | Coefficient of variation in age at reproduction. | $S = \sqrt{\bar{V}_f} / \bar{a}_f$ |
| Net reproductive rate | $R_0$ | Mean number of recruits produced during the mean life expectancy of an individual in the population. | $R_0 = \sum_{x=0}^{x=\eta_e} l_x m_x$ |
| Mature life expectancy | $L_\omega$ | Number of years from the mean age at maturity ($L_a$) until the mean life expectancy ($\eta_e$) of an individual in the population. | $L_\omega = \eta_{L_p}$ |

To identify life history strategies along major axes, we applied a varimax-rotated, mass corrected PCA using the *prcomp* function from the *stats* package in the statistical software R (R Core Team, 2023). The derived traits were log-transformed and scaled to adhere to PCA assumptions (*μ*= 0 and SD= 1; Lucas *et al.,* 2024).

The influence of phylogeny and body mass on the structuring of life history strategies was checked by running PCAs with and without correction for phylogeny, mass, or mass and phylogeny combined (Lucas et al. 2024). To correct for phylogenetic relatedness a species-level phylogenetic tree was constructed (Figure A1; Appendix) using data from the TimeTree database (Kumar et al. 2022). Phylogenetically corrected PCAs (pPCAs) were then calculated using the *phyl.pca* function from the package *phytools* (Revell 2012). The pPCA links the phylogenetic relationship of species to the life-history traits via a modified covariance matrix and estimated Pagel’s *λ* (Lucas et al. 2024). Pagel’s *λ* is a scaling parameter for the correlation in traits between species ranging from 0 (no correlation) to 1 (the correlation expected under Brownian motion), where any value greater than 0.5 suggests a somewhat influential role of overall phylogenetic ancestry (Freckleton et al. 2002). To correct for body mass the maximum female mass values for each species were gathered using targeted literature searches (Table 1). Body mass was then corrected for by computing the residuals for linear models between the log10-transformed derived life history traits (Table 2) and the body mass for each species (Table 1) for both the PCA and the pPCA (Revell 2009). The influence of these corrections was then assessed using Pagel’s *λ* and qualitative comparison of the PCA loadings to identify the most appropriate correction to apply for all further analyses (Lucas et al. 2024). Finally, the Kaiser criterion was applied to assess the significance of the PC axes, so that only the axes that were influential on the species life history strategy were retained (Lucas et al. 2024). The Kaiser criterion specifies that a PC axis is influential if it has an eigenvalue greater than one (Legendre & Legendre 2012).

### 2.2. Linking life history strategies to population performance and conservation (objective 2)

To test whether species’ position on the influential PC axes can predict variables of conservation interest (objective 2), we conducted a series of generalized linear models (GLMs) to test whether PC scores, and the two-way interactions between those scores, predicted a series of response variables of conservation interest (Salguero-Gómez 2017; Lucas et al. 2024): population growth rate, damping ratio (a measure of demographic resilience), elasticity (a measure of sensitivity to perturbation), and extinction risk.

When considering the value of these variables to conservation planners, a higher population growth rate indicates the number of individuals in the population will increase more rapidly than in a population with a lower population growth rate, which may improve the ability of a population to survive stochastic events or respond to change (Sutherland & Norris 2002).

Likewise, a larger damping ratio indicates a population will converge faster and recover in less time following a stochastic event or change (Stott et al. 2011; Capdevila et al. 2020). An elasticity analysis is one type of perturbation analysis, that estimates the sensitivity of species to a proportional change in each of the input life history parameters, by calculating the effect on the population growth rate (de Kroon et al. 1986). This can be used to pinpoint which life history parameter is most influential on a population’s growth rate and how this varies at different feeding levels, which in turn can be used to consider how conservation planners target interventions (Benton & Grant 1999; Rademaker et al. 2024).

To calculate population growth rate (λ), we discretised each DEB-IPM by dividing the length domain Ω (Equation 1) into 200 discrete bins, resulting in a matrix **A** of size *m* × *n*, where *m* = *n* = 200, and which dominant eigenvalue equals λ (Lucas et al. 2024). To calculate demographic resilience, we calculated the damping ratio (ξ), with ξ = λ/||λ_2_||, where λ_2_ is the highest subdominant eigenvalue of matrix A (Capdevila et al. 2020). We conducted an elasticity analyses for each species to examine which of the eight life history parameters (Table 1) most strongly affected their population growth rate (Smallegange & Lucas 2024). After deriving life history traits for each species across a range of feeding levels between 0.5 and 1.0, we used elasticity analysis to identify which life history trait was most influential at our chosen feeding level of 0.9 (Smallegange & Lucas 2024).

We conducted GLMs for all response variables, testing whether PC scores, and the two-way interactions between those scores predicted each response variable individually. All GLMs were conducted in R, and the model assumptions for each GLM were checked for homoscedasticity and collinearity <u>(Nelder & Wedderburn 1972; Fox et al. 2007; Hothorn et</u> <u>al. 2015; R Core Team 2023)</u>. The GLMs for population growth rate and demographic resilience were constructed as Gaussian models using the “identity” link function, and both GLMs were checked for normal distribution of residuals (Nelder & Wedderburn 1972).

Elasticity was treated as a binomial variable because the population growth rate was most sensitive to perturbation of only two life history traits: length at maturity (*L*_m_) and length at puberty (*L*_p_). Extinction risk (Table 1; IUCN, 2024) was also treated as a binomial variable; due to the small sample size there were not enough replicates in each individual extinction risk category and so species were classed as either threatened (vulnerable, endangered and critically endangered) or not threatened (least concern and near threatened). The GLMs for elasticity and extinction risk were constructed as Binomial models using the “logit” link function, and both GLMs were checked for overdispersion (Nelder & Wedderburn 1972). We used forward and backward stepwise selection to determine the best-fit model by comparing model AIC (Akaike 1998).

## 3. Results

### 3.1. Species’ life history strategies (objective 1)

When considered alone, phylogenetic relationships played a role in explaining the variety of life history strategies in the dataset. The Pagel’s *ʎ* value of the phylogenetic PCA was 0.56, suggesting a somewhat influential role of overall phylogenetic ancestry in our analyses (Table A1, Appendix; Freckleton, Harvey and Pagel, 2002). However, when the mass corrected PCAs with and without phylogenetic corrections were compared there was high qualitative similarity in the loadings of each axis, suggesting that, once the data had been corrected for mass, phylogeny did not change the results (Table A2, Appendix). Because of this, we continued with the mass corrected, non-phylogenetic PCA for all further analyses.

Following the Kaiser criterion (Legendre & Legendre 2012) PC axes 1, 2, 3 and 4 were retained in the analyses (Table A3, Appendix). The variety of life histories among reptiles was captured almost entirely by these four PC axes, which cumulatively explained 96% of the variation (Table 4). Following analysis, we only focus on the first three axes in the discussion as these were significant predictors and cumulatively explain 83% of the variation.

**Table 3:**
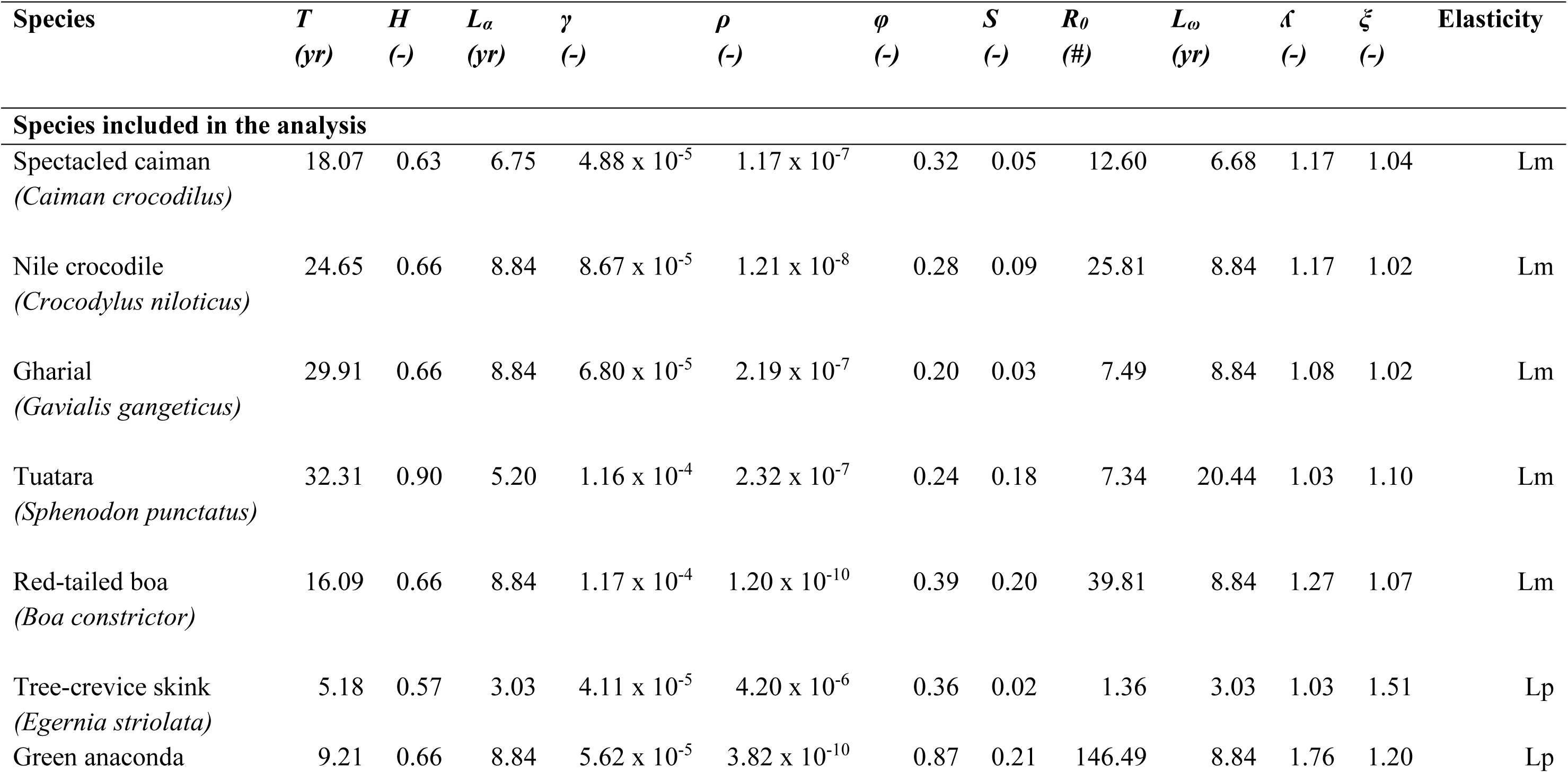
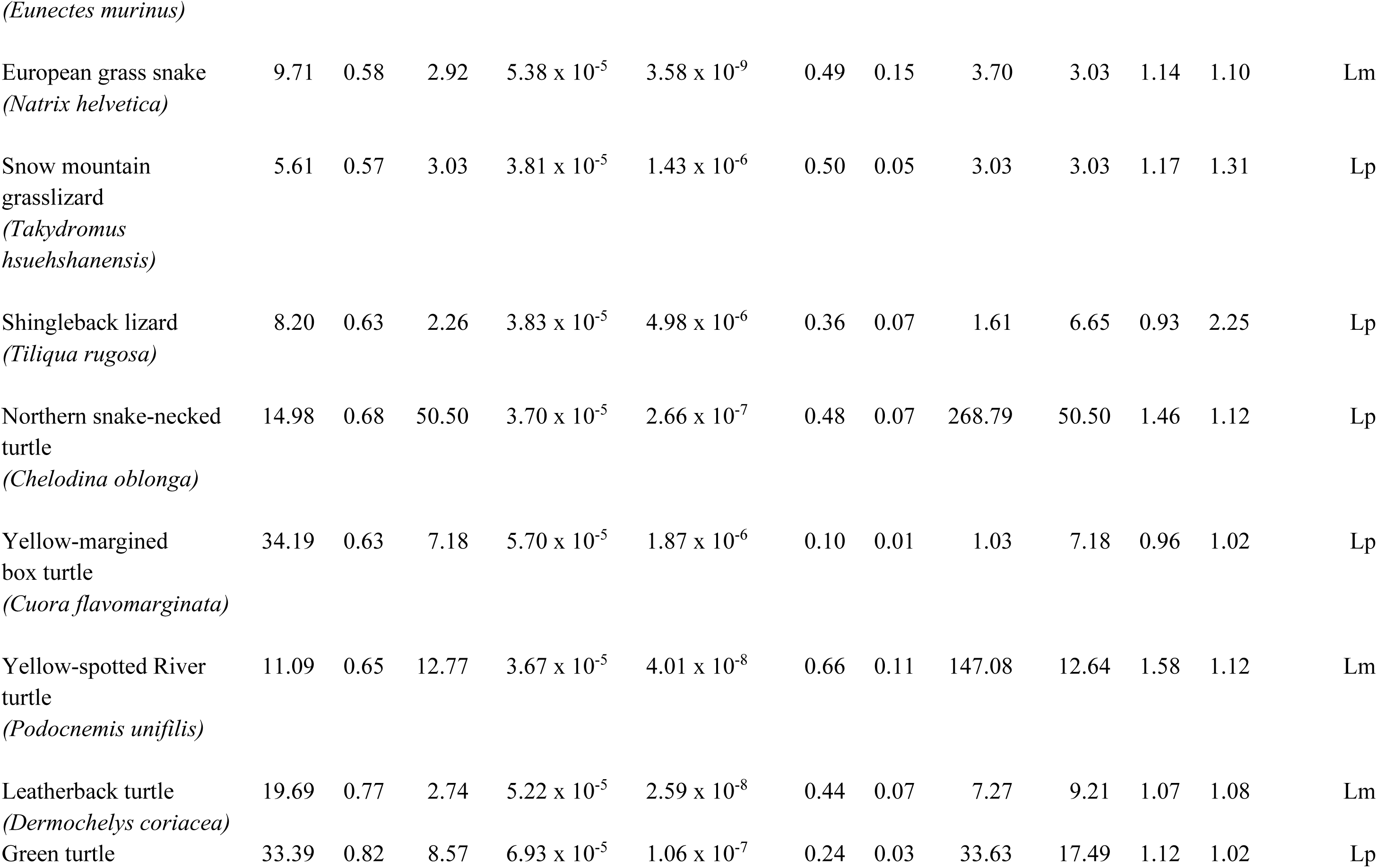
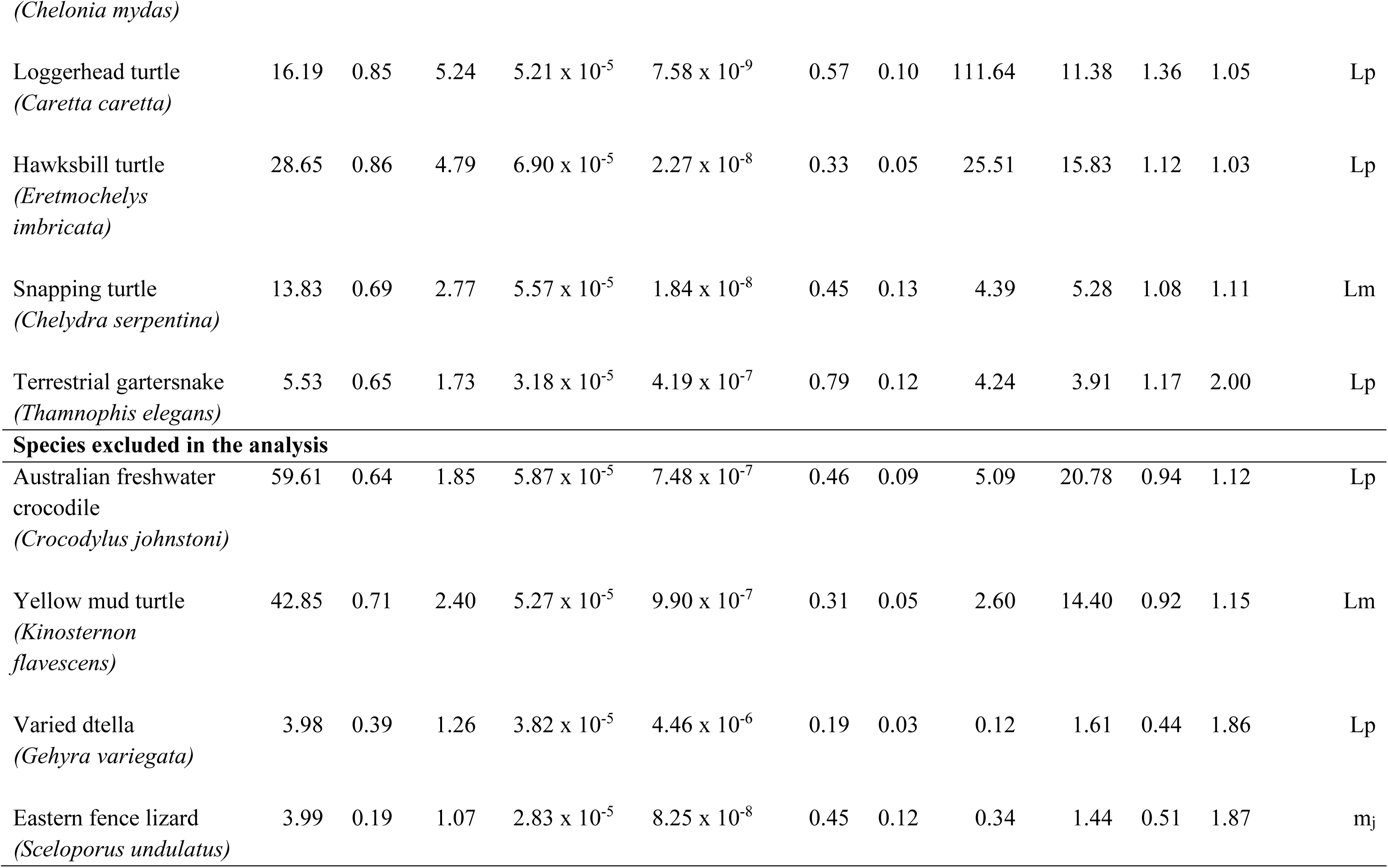
Derived life history trait values for each species at a feeding level of 0.9. T = generation time, H = survivorship curve, L*_α_* = age at maturity, γ = progressive growth, ρ = retrogressive growth, φ = mean recruitment success, S = degree of iteroparity, R_0_ = net reproductive rate, L_ω_ = mature life expectancy, ʎ = population growth rate and ξ = damping ratio. The elasticity column indicates which life history parameter was most influential on population growth rates at the given feeding level; for this dataset all species were either limited by length at puberty (*L_p_*), maximum length (*L_m_*) or juvenile mortality rate (μ*_j_*).

**Table 4:**
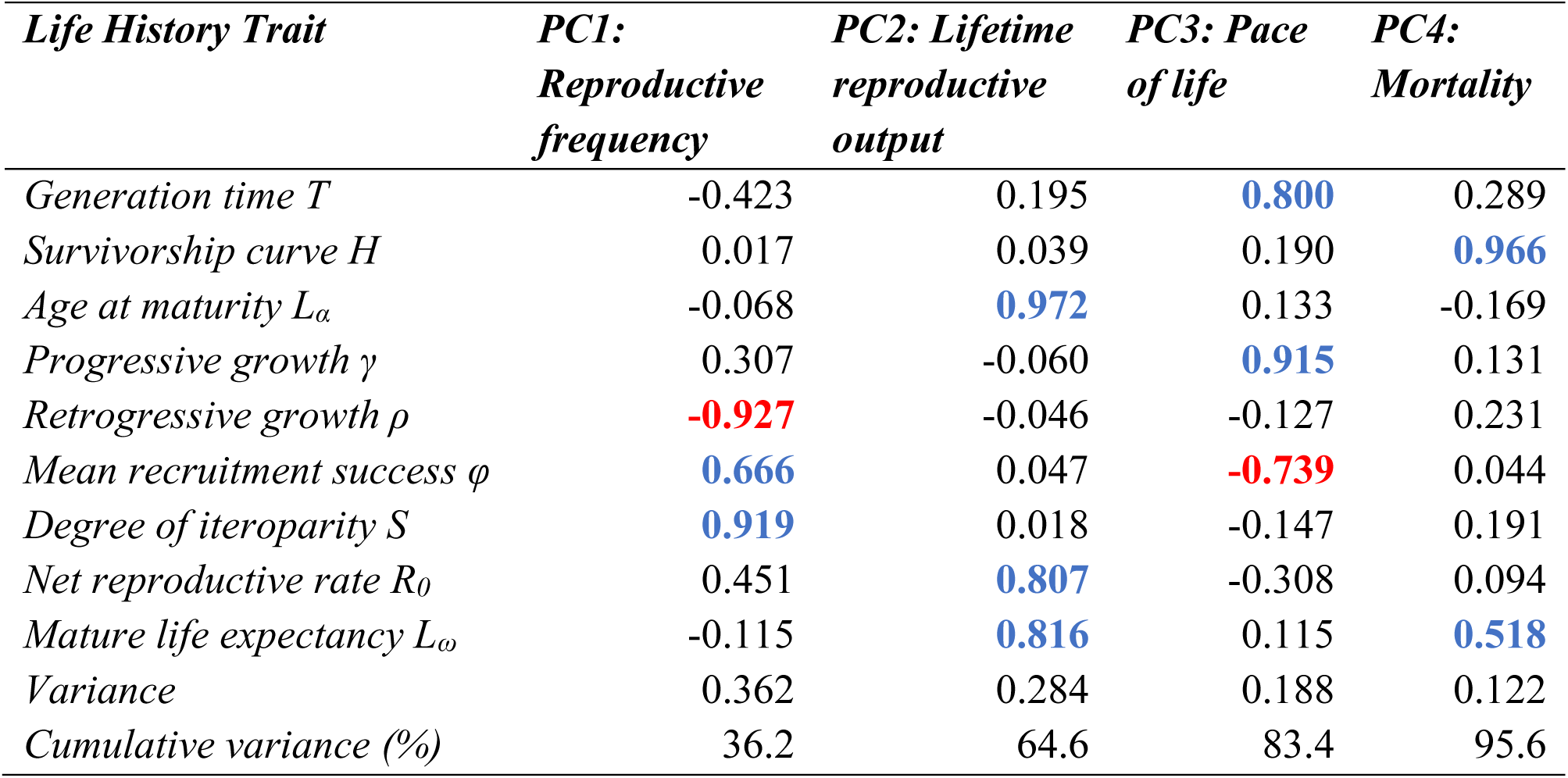
Loadings of the derived life history traits onto the first four axes of mass corrected PCA at a feeding level of 0.9. Loadings in bold indicate a high contribution (greater than ±0.50) to the given PC axis, with red indicating negative loadings while blue indicates positive.

| *Life History Trait* | *PC1:*<br>*Reproductive*<br>*frequency* | *PC2: Lifetime*<br>*reproductive*<br>*output* | *PC3: Pace*<br>*of life* | *PC4:*<br>*Mortality* |
| --- | --- | --- | --- | --- |
| *Generation time $T$* | -0.423 | 0.195 | **0.800** | 0.289 |
| *Survivorship curve $H$* | 0.017 | 0.039 | 0.190 | **0.966** |
| *Age at maturity $L_a$* | -0.068 | **0.972** | 0.133 | -0.169 |
| *Progressive growth $\gamma$* | 0.307 | -0.060 | **0.915** | 0.131 |
| *Retrogressive growth $\rho$* | **-0.927** | -0.046 | -0.127 | 0.231 |
| *Mean recruitment success $\phi$* | **0.666** | 0.047 | **-0.739** | 0.044 |
| *Degree of iteroparity $S$* | **0.919** | 0.018 | -0.147 | 0.191 |
| *Net reproductive rate $R_0$* | 0.451 | **0.807** | -0.308 | 0.094 |
| *Mature life expectancy $L_\omega$* | -0.115 | **0.816** | 0.115 | **0.518** |
| *Variance* | 0.362 | 0.284 | 0.188 | 0.122 |
| *Cumulative variance (%)* | 36.2 | 64.6 | 83.4 | 95.6 |

The life history traits most closely aligned with PC axes 1 and 2 were both related to reproductive strategy: retrogressive growth (*ρ*) was the life history trait with the greatest loading onto PC axis 1 (Table 4), followed closely by degree of iteroparity (*S*) and then mean recruitment success (*φ*). To simplify interpretation, this axis was inverted, so that both S and *φ* were positively loaded onto the axis, while *ρ* had a negative loading, showing an inverse association between species reproductive output and the likelihood of a species undergoing retrogressive growth in poor environments. The trait with the greatest loading onto PC axis 2 was age at maturity (*L_α_*), followed by mature life expectancy (*L_ω_*) and net reproductive rate (*R_0_*). All three traits were positively loaded onto the axis, suggesting species with a longer life expectancy had greater net reproductive rates.

The life history traits most closely aligned with PC axis 3 were related to the fast-slow continuum (Gaillard et al. 2016): the trait with the greatest loading was progressive growth (*γ*), followed by generation time (*T*). Both were positively loaded onto PC axis 3 while mean recruitment success (*φ*) was negatively loaded onto the axis (Table 4), suggesting an inverse association between reproduction and generation time, while generation time had a positive association with growth. The life history traits most closely aligned with PC axis 4 were related to species mortality: the trait with the greatest loading onto PC axis 4 was the shape of the survivorship curve (*H*), followed by mature life expectancy (*L_ω_*), and both were positively loaded.

As scores increased along PC axis 1 (hereafter the reproductive frequency axis), reptiles exhibited increasing frequency of reproduction and recruitment success and decreasing tendency to shrink (Figure 2). As scores increased along PC axis 2 (hereafter the lifetime reproductive output axis), reptiles exhibited increasing allocation to longevity related life-history traits and net reproductive rate. As scores increased along PC axis 3 (hereafter the pace of life axis), reptiles exhibited increasing allocation to growth and generation time, and decreasing recruitment success. Finally, as scores increased along PC axis 4 (hereafter the mortality axis), species exhibited increasing their allocation to longevity related life history traits and decreasing mortality with age.

**Figure 2:**
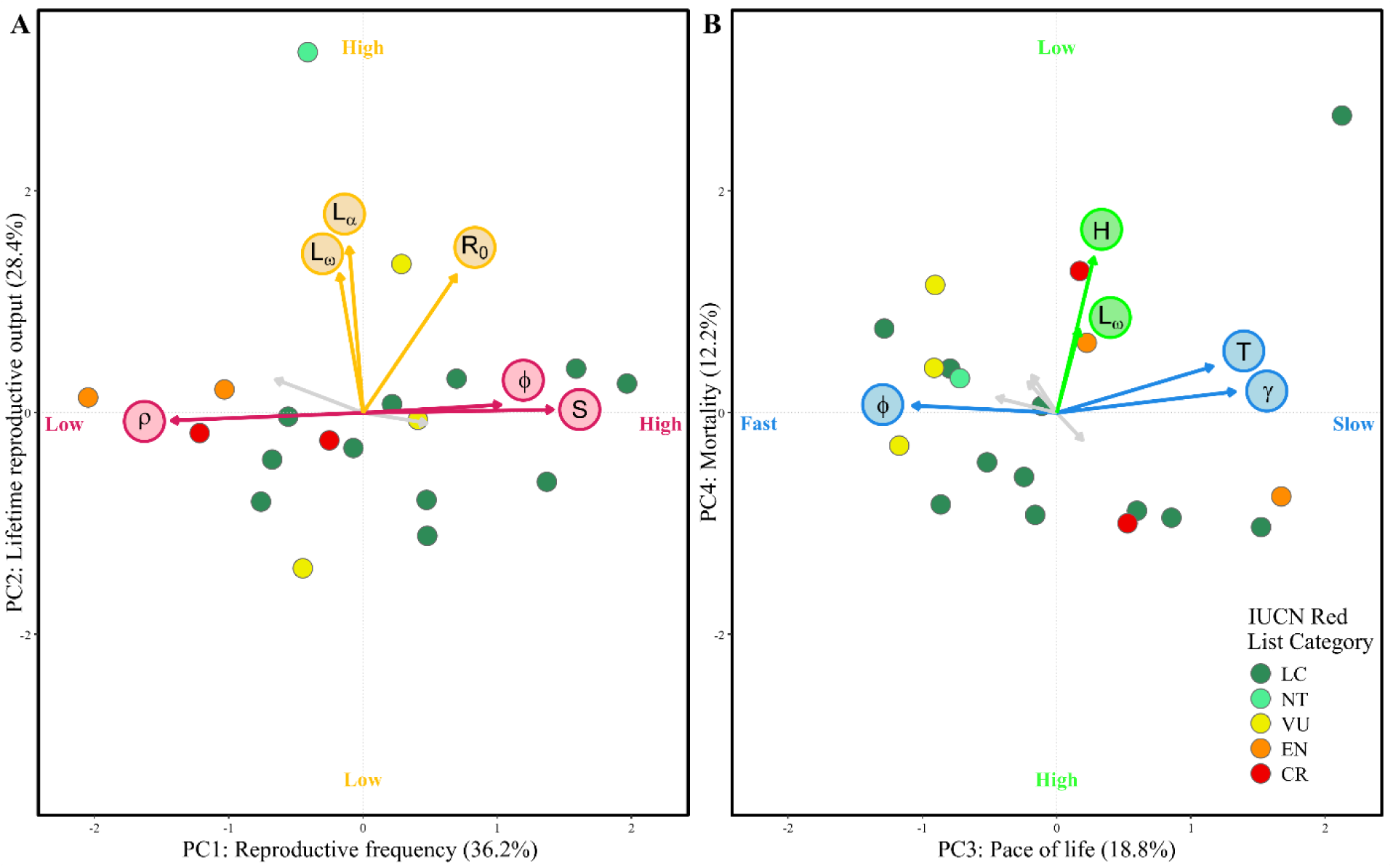
Mass corrected PCA of life history strategies showing loadings of each derived life history trait. (A) PC axes 1 (reproductive frequency) and 2 (lifetime reproductive output), with the loadings of PC1 in pink and PC2 in yellow. (B) PC axes 3 (pace of life) and 4 (mortality) with the loadings of PC3 in blue and PC4 in green. The points are coloured according to the species extinction risk on the IUCN Red List. Percentages in brackets shows the percentage variance in life histories explained by the PC axis.

### 3.2. Life history strategies as predictors of variables of conservation interest (objective 2)

The position of a species along our life history strategies axes was found to predict its demographic performance. The population growth rate of species increased significantly as the species ranked higher on scores along both the reproductive frequency axis (i.e. reproduced more frequently and higher recruitment; Figure 3; Table 5), and the lifetime reproductive output axis (i.e., higher longevity and net reproductive rate; Table 5).

**Figure 3:**
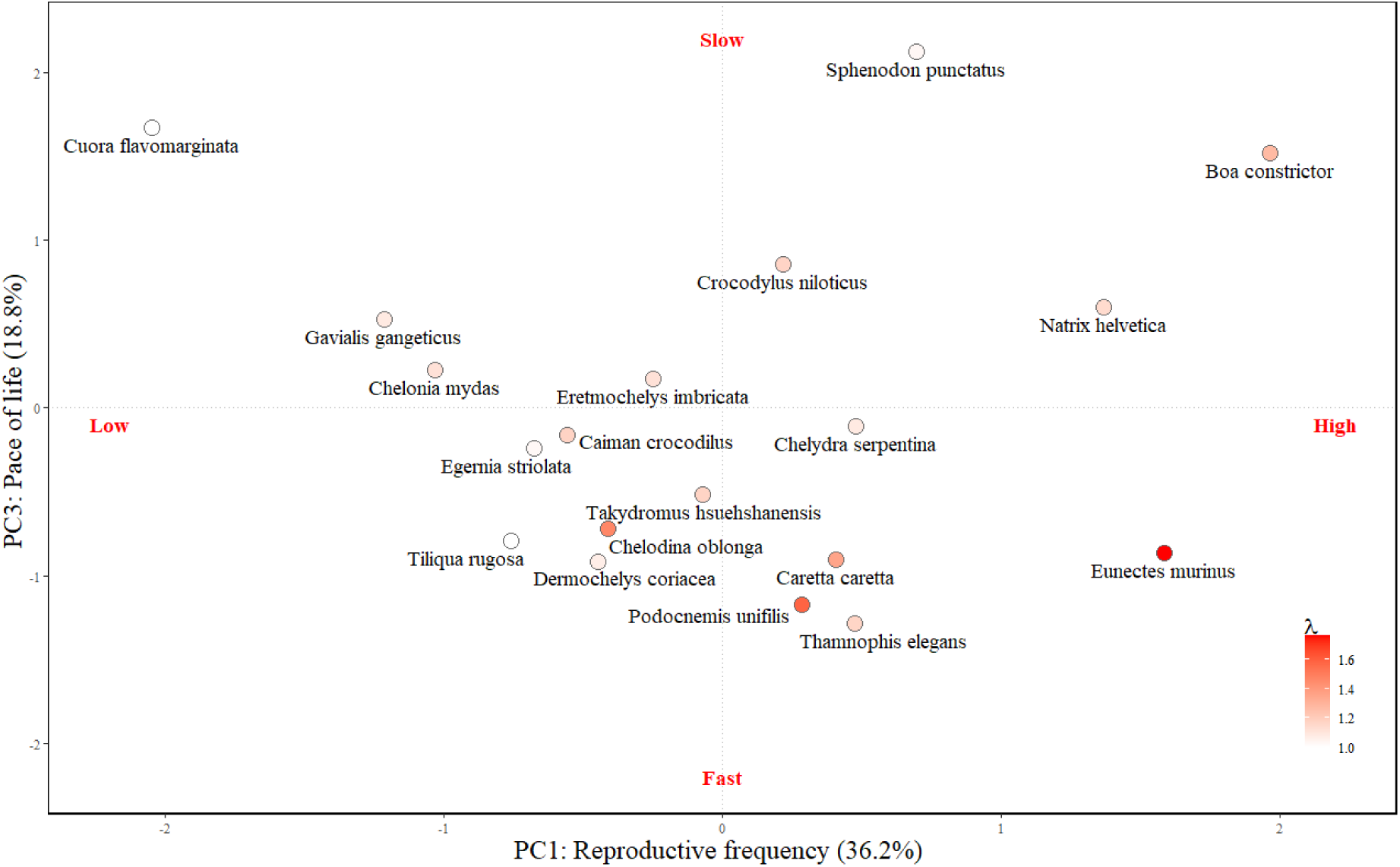
Species position along PC axes 1 (reproductive frequency) and 3 (pace of life), which were significant predictors of population growth rate (Table 5). Points are coloured by population growth rate. Species labels are all underneath their associated point. Percentages in brackets show the percentage of variance in life histories explained by the PC axis.

**Table 5:** Generalised linear models for population growth rate (*ʎ*), demographic resilience (ξ), IUCN threat status and elasticity, as a response to PC1 (reproductive frequency), PC2 (lifetime reproductive output), PC3 (pace of life), PC4 (mortality) and the two-way interactions between these predictor variables. Both IUCN threat status and elasticity were modelled as binomial response variables. Models were refined using forward and backward stepwise selection to determine the best-fit model by comparing model AIC (Akaike 1998).

| **Population growth rate ($\lambda$)** |  |  |  |  |
| --- | --- | --- | --- | --- |
| **Predictor** | **Coefficient**<br>**(Estimate)** | **Standard error** | **t-value** | **p-value** |
| (Intercept) | 1.194 | 0.013 | 91.037 | >0.001 |
| PC1 | 0.154 | 0.017 | 9.293 | >0.001 |
| PC2 | 0.143 | 0.015 | 9.759 | >0.001 |
| PC3 | -0.073 | 0.014 | -5.272 | >0.001 |
| PC4 | -0.021 | 0.014 | -1.536 | 0.150 |
| PC1:PC2 | 0.104 | 0.029 | 3.644 | 0.003 |
| PC1:PC3 | -0.072 | 0.014 | -5.150 | >0.001 |
| **Demographic resilience ($\xi$)** |  |  |  |  |
| **Predictor** | **Coefficient**<br>**(Estimate)** | **Standard error** | **t-value** | **p-value** |
| (Intercept) | 1.221 | 0.069 | 17.776 | >0.001 |
| PC2 | 0.100 | 0.145 | 0.687 | 0.503 |
| PC3 | -0.191 | 0.076 | -2.504 | 0.024 |
| PC2:PC3 | 0.280 | 0.181 | 1.548 | 0.143 |
| **IUCN Threat status** |  |  |  |  |
| **Predictor** | **Coefficient**<br>**(Estimate)** | **Standard error** | **z-value** | **p-value** |
| (Intercept) | -7.461 | 6.230 | -1.198 | 0.231 |
| PC1 | -8.752 | 5.956 | -1.469 | 0.142 |
| PC2 | -8.481 | 7.782 | -1.090 | 0.276 |
| PC3 | -4.848 | 4.667 | -1.039 | 0.299 |
| PC4 | 4.965 | 3.174 | 1.565 | 0.118 |
| PC1:PC3 | -7.724 | 5.650 | -1.367 | 0.172 |
| PC2:PC3 | -10.246 | 8.157 | -1.256 | 0.209 |
| **Elasticity** |  |  |  |  |
| **Predictor** | **Coefficient**<br>**(Estimate)** | **Standard error** | **z-value** | **p-value** |
| (Intercept) | -0.417 | 0.775 | -0.539 | 0.590 |
| PC1 | -0.947 | 0.978 | -0.968 | 0.333 |
| PC3 | -1.354 | 1.025 | -1.321 | 0.186 |
| PC4 | 1.639 | 1.063 | 1.542 | 0.123 |
| PC1:PC3 | -2.086 | 1.473 | -1.416 | 0.157 |

Additionally, as species scored higher on the pace of life axis (i.e., lower recruitment but higher longevity and growth rate), their population growth rates declined significantly (Figure 3; Table 5). The interactions between species position on the reproductive frequency and lifetime reproductive output axes, and on the reproductive frequency and pace of life axes were also significant (Figure 3; Table 5). In other words, we would expect species with high reproductive outputs and fast life history strategies, that have a shorter lifespan but grow more quickly, to show higher population growth rates following successful conservation intervention. A species position on the mortality axis did not significantly predict population growth rates.

As a species scored higher along the pace of life axis (i.e., lower recruitment but higher longevity and growth rate), their demographic resilience decreased significantly (Table 5; Figure 4). Thus, we would expect species with fast life history strategies, that have a shorter lifespan but produce more offspring and grow more quickly, to be more sensitive to and less able to recover from disturbances. However, the species position on the reproductive frequency, lifetime reproductive output and mortality axes did not significantly predict demographic resilience (Table 5). The position of species on life history strategy axes did not significantly influence the extinction risk of species or sensitivity to perturbations (Table 5; Figure A2, Appendix).

**Figure 4:**
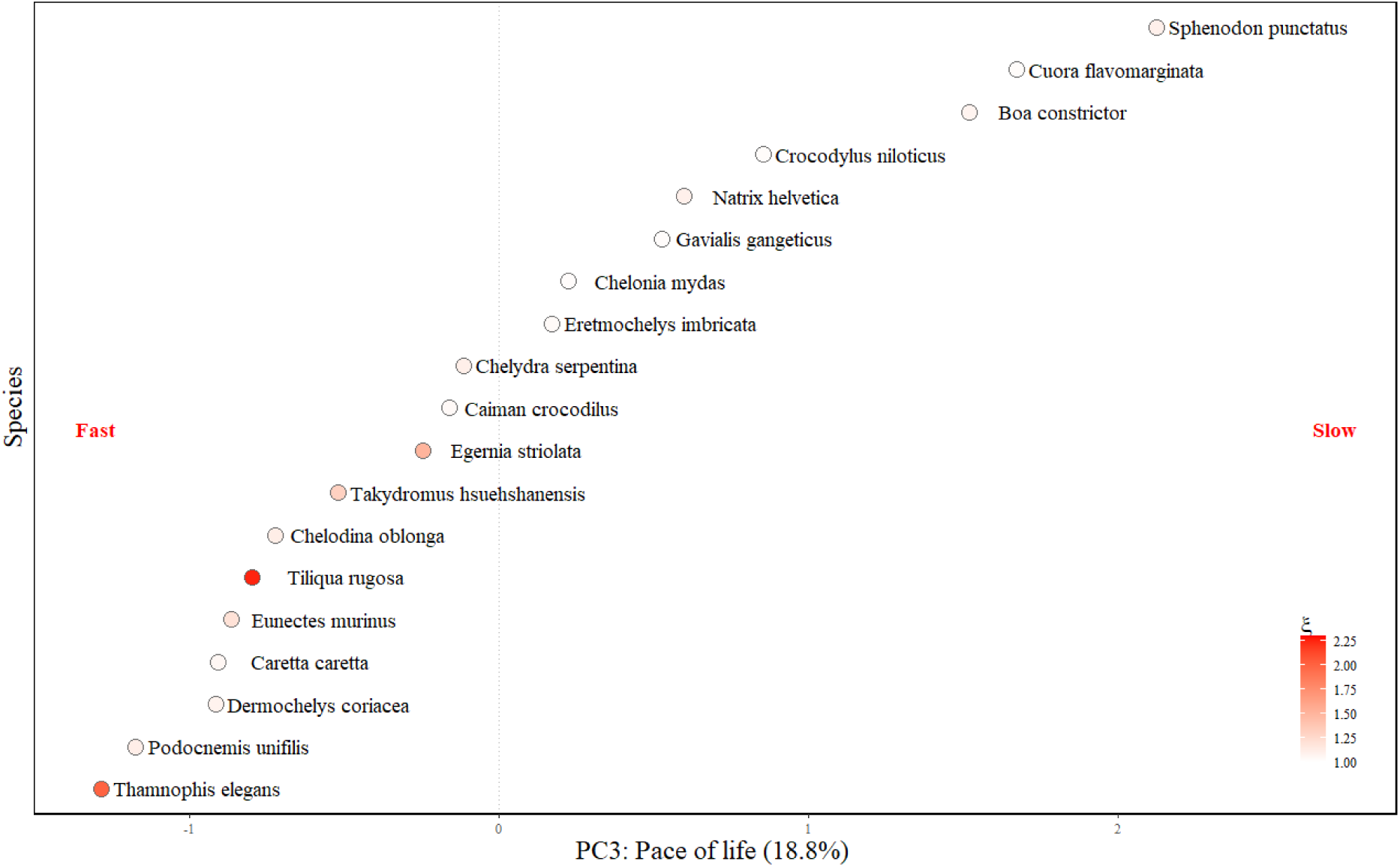
Species position along PC axis 3 (pace of life), which was a significant predictor of demographic resilience. Points are coloured by demographic resilience. Percentages in brackets show the percentage of variance in life histories explained by the PC axis.

## 4. Discussion

Our study investigated the extent to which life history strategies predicted variables of conservation interest for reptiles to assess whether life history strategies have the potential to inform conservation intervention planning. Using a representative selection of species, including all extant orders and a mixture of threatened and non-threatened species, we found that reptile life history strategies were structured along four independent axes, corresponding to reproductive frequency, lifetime reproductive output, pace of life and mortality (objective 1). A species position on the reproductive frequency, lifetime reproductive output and pace of life axes predicted population growth rate, suggesting we can use life history data to inform our understanding of reptile species conservation recovery potential, by predicting the likely population growth rate after conservation interventions. A species position along the pace of life axis also predicted the populations’ demographic resilience, informing our understanding of species potential for recovery after conservation interventions or disturbance (objective 2).

### 4.1. Informing conservation action

We found that the population growth rate of reptiles was significantly influenced by their position along the reproductive frequency, lifetime reproductive output and pace of life axes; fast-lived species with high reproductive frequency and output exhibited higher population growth rates. Estimates of population growth rates are valuable to conservation planners as they could be used to predict the likely population growth rate after conservation interventions (Piipponen-Doyle et al. 2021). In cases where there is limited capacity or budget, deciding the most efficient period to conduct monitoring following a conservation intervention can be influenced by the predicted time needed to show a significant change in the population (Grainger et al. 2019; Piipponen-Doyle et al. 2021). Where species are predicted to have a lower population growth rate based on their life history strategies, conservation planners should consider the need to monitor for a long period of time – the cost of which could be partially offset by long intervals between monitoring efforts – in order to evidence the impact of interventions. Similarly, by applying these population predictions it may be possible to determine that conservation interventions have not been successfully implemented, or may have targeted the wrong threat, by demonstrating that species populations have not recovered as expected (Bakker & Doak 2009). By gathering conclusive evidence of what works, and ensuring this information is shared, this can support future conservation planning, and prevent wasting time and resources (Sutherland et al. 2004; Catalano et al. 2019). In turn this evidence could support efforts to prove interventions have reduced the extinction risk of species as part of Goal A of the KMGBF (CBD, 2022).

We also found that a species’ position along the pace of life axis was able to predict the populations’ demographic resilience, suggesting that fast-lived reptiles exhibit increased demographic resilience to change and higher potential for recovery (Stott et al. 2011). Using this measure of demographic resilience could help conservation planners by indicating the time needed for a population to return to a stable age distribution, reducing the uncertainty about how a species is expected to recover following conservation interventions (Watts et al. 2020; Capdevila et al. 2020). This information could also be used to prioritise conservation interventions among threatened species, for example if a species with limited distribution that is threatened by stochastic events (e.g. an island endemic) is shown to have low demographic resilience this could support conservation planners to prioritise ex-situ conservation or establish other populations (McGowan et al. 2017). Within Goal A of the KMGBF there is also the agreement that “the abundance of native wild species is increased to healthy and resilient levels” (CBD, 2022). This approach therefore has direct value to conservation planners and policy makers as a method to generate a metric of demographic resilience for data-sparse species without extensive data gathering (Smallegange et al. 2017).

Our elasticity analysis showed that the position of reptile species on our life history strategy axes did not predict which life history parameter would be most sensitive to perturbations. This lack of significant relationship may be because our populations were modelled at a feeding level of 0.9, which allowed us to maximise our sample size but represents a “well fed” population (Smallegange et al. 2017). Different results may be obtained for lower feeding levels, as life history traits were shown to vary when calculated at different feeding levels (Lucas et al. 2024). Elasticity analyses can nevertheless be very useful when planning conservation interventions, by preventing prioritisation of efforts that may not be as effective at reducing extinction risk: studies on sea turtles have demonstrated the importance of prioritising adult survival over the more widely implemented intervention of increasing small juvenile survival using methods such as protecting eggs in artificial beach hatcheries (Crowder et al. 1994; Richards et al. 2024).

We also found no significant relationship between the position of reptile species on our life history strategy axes and their extinction risk. These results differ from the findings of Lucas *et al.,* (2024) and Salguero-Gómez (2017), where slow-lived elasmobranch and plant species were shown to be significantly more likely to be classified as threatened on the IUCN Red List (Salguero-Gómez 2017; Lucas et al. 2024). This may be a result of the high variation in lifespan exhibited by species within both those groups, with some plant species recorded to have a maximum longevity of thousands of years, making them disproportionately vulnerable to certain forms of disturbance (Piovesan & Biondi 2021). However, our results do support the findings of Healy *et al.,* (2019), who found no significant relationship between species position on a life history strategy framework and extinction risk when a broad variety of animal species were compared. Our finding is still important for conservation planners, as it suggests that all reptiles species should be considered at risk of becoming threatened, with no single life history strategy being less susceptible than another to the threats species currently face (Healy et al. 2019; Finn et al. 2023).

### 4.2. Life history framework in reptiles

Our study focused on three influential axes of life history strategies, which explained 84% of the variation in life history strategies, possibly reflecting the variation in morphology, physiology, and behaviour among reptile orders (Hallmann & Griebeler 2018; Taylor et al. 2021). Studies of other taxonomic groups have found two influential axes (Salguero-Gómez et al. 2016; Healy et al. 2019), and our results support growing evidence that reptile life-history strategies do not directly align with other classes of species (Healy et al. 2019; Wright et al. 2020). For example, lizards of the Lacertidae family do not follow the strategies described by the fast-slow continuum, with some species having small clutches but relatively large young, placing them at the “fast” end of the continuum, and others having large clutches and small young, placing them at the “slow” end (Bauwens & Diaz-Uriarte 1997).

Our finding of two influential reproductive axes may reflect the wide variety in reproductive strategies exhibited by reptiles; some species are semelparous (i.e., reproduce once in a lifetime) while others reproduce annually over many years, and some species consistently produce the same clutch size while others can differ by dozens (Shine & Greer 1991).

Reproduction also varies from “live-birth” (viviparity) to “egg-laying” (oviparity; Shine, 1983), which has been described as the oviparity-viviparity continuum. Along this continuum species also exhibit variation in oviparity, with some retaining eggs in utero for only a brief time while other retain eggs in utero for most of the embryonic development (Shine 1983).

Our results demonstrate the importance of studies focusing within taxonomic groups alongside broad cross-taxonomic comparisons, to avoid missing taxon-specific insights (Rademaker et al. 2024).

We also found that body mass and phylogeny were associated for our sample of reptiles. Variation in body mass among the Squamata and Rhynchocephalia orders has been found to span six orders of magnitude, with the mean weight of snakes being seven times heavier than lizards (Feldman et al. 2016). Another study on squamate reptiles showed that large-bodied species lived longer than small ones, with brood frequency negatively correlating with longevity (Scharf et al. 2015). Our pace of life axis demonstrated the same relationship, where generation time was positively associated with growth but negatively associated with mean recruitment success. Although some studies show the same relationship between life history traits in other taxon (Lucas et al. 2024), this differs from the expected relationship, where investment in growth is associated with species with faster life histories, as for plants (Salguero-Gómez et al. 2016). These differences are likely because of reptile growth strategies as – unlike many species of birds, mammals and plants – many reptiles continue to grow after reaching sexual maturity (Hughes 2017; Hallmann & Griebeler 2018).

### 4.3. Study Constraints

Individual-level models allow for greater complexity but typically examine a smaller sample of species, which is contrasted against broad cross-taxonomic comparisons that can use large functional trait datasets, allowing increased statistical power, but at the cost of reduced mechanistic insight (Rademaker et al. 2024). The IUCN Red List has assessed 10,309 species of reptiles (IUCN, 2024), of which over 96% belong to the Squamata order (Roll et al. 2017). In our study we represented all the orders within Reptilia, with nine species from the Squamata order, nine Testudines, four Crocodilia and the single species belonging to Rhynchocephalia. To increase representation in future, the dataset could be expanded to comprise a greater proportion of squamate species.

As part of data gathering, we collected body mass data from published literature. While methods often described collection, the resulting body mass data were not always published (see Gregory, Isaac and Griffiths, 2007, for an example from our data sources). Body mass can be calculated using body length (e.g. Feldman *et al*., 2016), but this approach has its own limitations (Campione & Evans 2012). Given the challenges associated with gathering data from fully grown individuals, both from a logistical and safety standpoint (Gienger et al. 2017), it is important that the data that have already been gathered are made publicly available (Reichman et al. 2011).

## 5. Conclusions and future research considerations

In this study we showed that life history data can be used to predict the population performance and demographic resilience of a representative selection of reptile species. This information has direct value to conservation planners, supporting informed predictions of how species will respond to conservation interventions in species with limited long-term population monitoring. The expansion of online open access databases are making life history trait data more easily accessible (Smallegange & Lucas 2024), which supports the potential for such data to be used in conservation decision making. Future research could test the predictions made using life history modelling against observed data, which would provide evidence for the reliability of life history analysis and encourage uptake of such analysis as an efficient conservation decision support tool.

## Author Contributions

Emily A. Stevenson: conceptualisation (equal), data curation (supporting), formal analysis (lead), writing – original draft (lead), writing – review and editing (equal).

Sol Lucas: data curation (supporting), formal analysis (supporting), methodology (supporting), writing – review and editing (equal).

Philip J. K. McGowan: conceptualisation (supporting), writing – review and editing (equal).

Isabel M Smallegange: conceptualisation (equal), data curation (lead), formal analysis (supporting), methodology (lead), writing – review and editing (equal).

Louise Mair: conceptualisation (equal), formal analysis (supporting), writing – original draft (supporting), writing – review and editing (equal).

## Acknowledgements

We thank Simon Maddock for contributing his expertise on reptiles. ES and LM were supported by funding from Newcastle University. SL was funded by the Leverhulme Doctoral Scholarship in Behaviour Informatics (DS-2017-015).

## Data accessibility statement

The data and MatLab code supporting this study were published by Smallegange and Lucas, (2024), and are available on FigShare (https://doi.org/10.6084/m9.figshare.13241972.v18).

## Appendix

## General structure of a DEB-IPM

A DEB-IPM is a population model that tracks cohorts of female individuals in a population, capturing the demographic rates of survival, growth, reproduction and the relationship between parent and offspring size, using eight life history parameters (Smallegange et al. 2017; Smallegange & Lucas 2024). By applying these demographic rates, studies have been able to predict the population performance, demographic resilience and extinction risk of other taxa, which could be utilised to support conservation planning and reduce uncertainty about species responses to conservation interventions (Lucas et al. 2024; Rademaker et al. 2024). Full details of how to parameterise and apply DEB-IPMs are in Smallegange and Lucas (2024); here, a brief description follows:

The number of females at year *t* is denoted by *N(L, t)* which allows the dynamics of body length number distribution from year *t* to *t* + 1 to be written as:

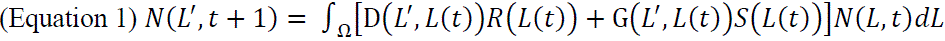

Where the closed interval Ω denotes the length domain (Smallegange et al. 2017).

The survival function *S*(*L*(*t*)) in equation (1) describes the probability that an individual of length *L* survives from time *t* to *t* + 1:

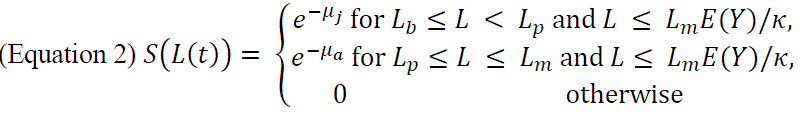

where *L_b_* is length at birth, *L_p_* is length at puberty and *L_m_* is the maximum attainable length (Table 1). Feeding level *E(Y)* can range from zero (empty gut) to one (full gut), and individuals die from starvation at a body length at which maintenance requirements exceed the total amount of assimilated energy (Smallegange et al. 2017). Starvation occurs when *L* > *Lm* · *E*(*Y*)/κ making *S*(*L*(*t*)) = 0 (i.e. an individual of length *L_m_* will die of starvation if *E(Y*) < κ, where κ is the fraction of assimilated energy allocated to respiration (Table 1), with 1 – κ allocated to reproduction). Juveniles and adults often have different mortality rates, and thus, juveniles (*L*_*b*_ ≤ *L* < *L*_*p*_) that do not die of starvation (i. e. *L* ≤ *Lm* · *E*(*Y*)/κ) have a mortality rate of *μ_j_*, and adults (*L*_*p*_ ≤ *L* ≤ *L*_*m*_) that do not die of starvation have a mortality rate of *μ_a_* (Smallegange & Lucas 2024).

The growth function *G*(*L*’, *L*(*t*)) in equation (1) describes the probability that an individual of length *L* at time *t* grows to length *L’* at time t + 1, conditional on survival, following a Gaussian distribution:

(Equation 3) *G*(*L*’, *L*(*t*)) =

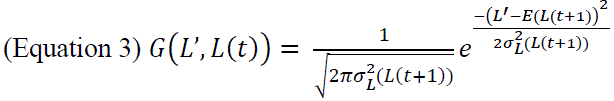

with the growth realised by a cohort of individuals with length *L*(*t*) equalling:

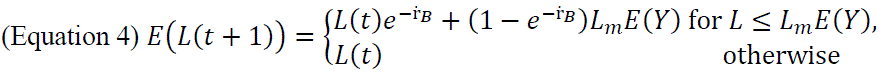

and the variance in length a time *t +* 1 for a cohort of individuals of length *L* as:

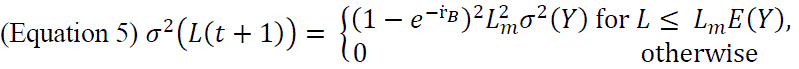

where σ(*Y*) is the standard deviation of the expected feeding level, and ṙ_*B*_ is the von Bertalanffy growth rate (Table 1; Smallegange *et al*., 2017). Individuals do not necessarily experience the same feeding level due to demographic stochasticity, which generates individual variability in the DEB-IPM (Smallegange et al. 2017).

The reproduction function *R*(*L*(*t*)) in equation (1) describes the number of offspring produced from time *t* to *t +* 1 by a female of length *L* at time *t*:

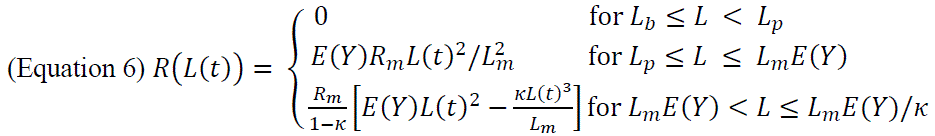

Individuals are mature when they reach puberty at body length *L_p_* and only surviving adults reproduce, meaning only individuals within a cohort of length *L*_*p*_ ≤ *L* ≤ *L*_*m*_*Y*/κ reproduce (Smallegange et al. 2017).

Finally, the probability density function, or parent-offspring association function, *D*(*L*’, *L*(*t*)) in equation (1) describes the probability that a female of length *L* at time *t* producing offspring of length *L’* at *t +* 1, conditional on reproduction, and hence describes the association between parent and offspring character values:

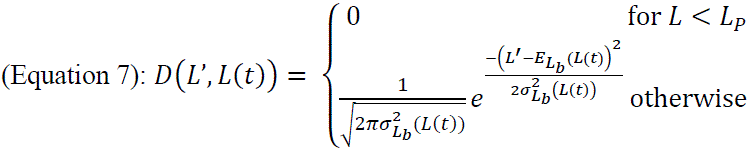

where 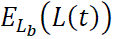 is the expected size of offspring produced by a cohort of individuals with length *L*(*t*), and 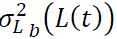 is the associated variance (Smallegange et al. 2017). For simplicity, 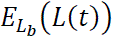 is set as a constant, and assume the variance 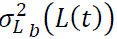 to be very small (Smallegange et al. 2017).

**Figure A1:**
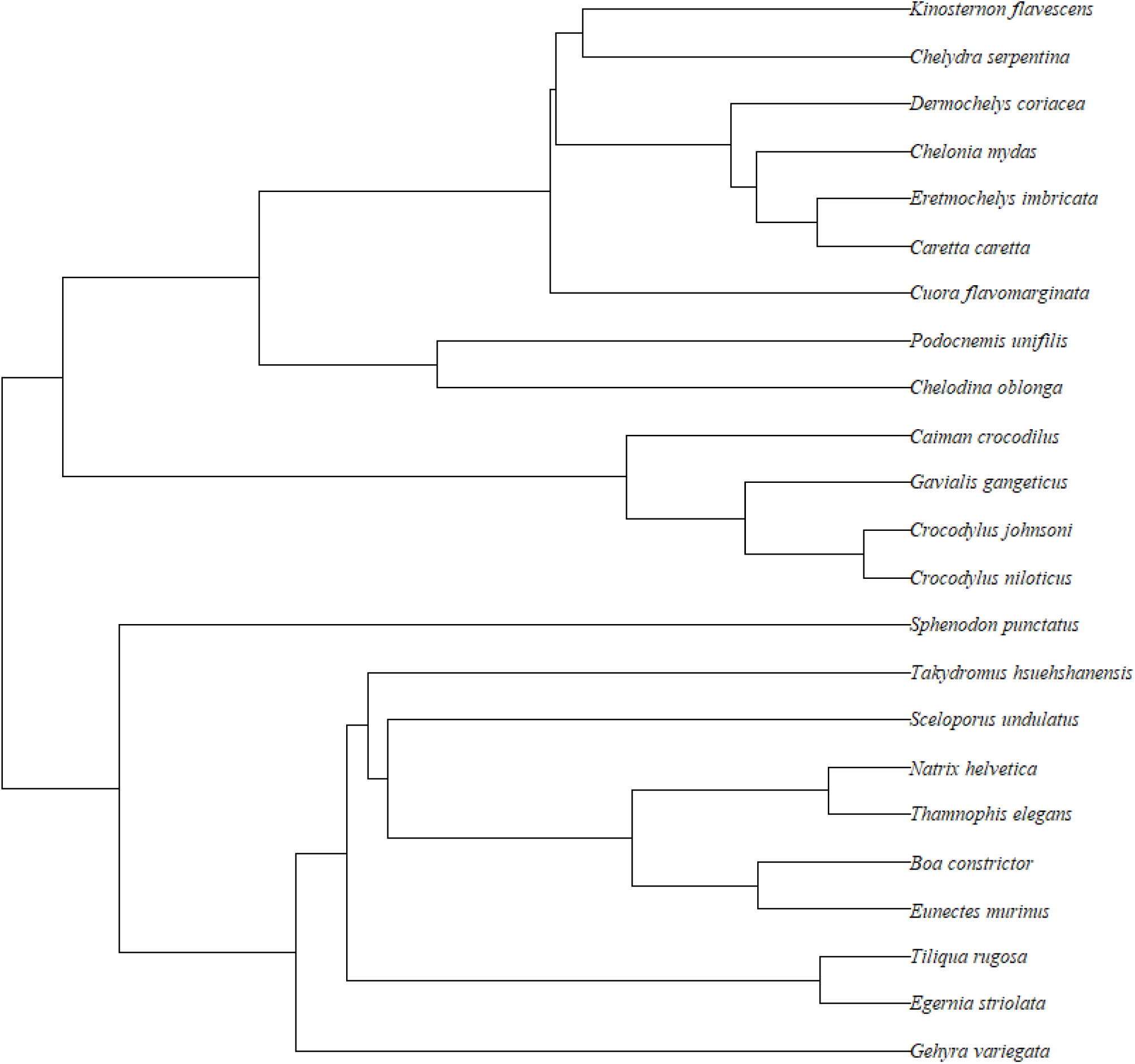
Phylogenetic tree of all reptiles on the DEBBIES database (Smallegange and Lucas, 2024), generated using the TimeTree database (Kumar et al., 2022).

**Table A1:** PCA loadings at feeding levels of 0.8 and 0.9 for non-phylogenetic and phylogenetically informed PCAs.

| Life History Trait | Feeding Level $E(Y)$ | Non-phylogenetic PCA | | | Phylogenetic PCA | | |
| --- | --- | --- | --- | --- | --- | --- | --- |
|  |  | PC1 | PC2 | PC3 | PC1 | PC2 | PC3 |
| Generation time $T$ | 0.8 | 0.525 | 0.130 | -0.121 | 0.911 | -0.177 | 0.292 |
| Survivorship curve $H$ | 0.8 | 0.491 | -0.070 | -0.011 | 0.838 | 0.106 | 0.152 |
| Age at maturity $L_a$ | 0.8 | -0.041 | 0.064 | -0.622 | 0.084 | 0.044 | 0.943 |
| Progressive growth $\gamma$ | 0.8 | 0.530 | -0.231 | 0.095 | 0.886 | 0.328 | 0.027 |
| Retrogressive growth $\rho$ | 0.8 | -0.165 | 0.541 | 0.019 | -0.317 | -0.831 | -0.174 |
| Mean recruitment success $\phi$ | 0.8 | -0.355 | -0.461 | -0.108 | -0.545 | 0.733 | 0.170 |
| Degree of iteroparity $S$ | 0.8 | 0.042 | -0.598 | 0.072 | 0.089 | 0.899 | 0.015 |
| Net reproductive rate $R_0$ | 0.8 | -0.096 | -0.212 | -0.555 | -0.009 | 0.452 | 0.880 |
| Mature life expectancy $L_\omega$ | 0.8 | 0.191 | 0.114 | -0.514 | 0.447 | -0.060 | 0.823 |
| Variance | 0.8 | 0.418 | 0.275 | 0.178 | 0.418 | 0.275 | 0.178 |
| Generation time $T$ | 0.9 | 0.517 | -0.156 | 0.118 | -0.181 | 0.916 | 0.271 |
| Survivorship curve $H$ | 0.9 | 0.399 | 0.113 | 0.104 | 0.420 | 0.534 | 0.256 |
| Age at maturity $L_a$ | 0.9 | -0.030 | -0.121 | 0.608 | -0.067 | 0.119 | 0.929 |
| Progressive growth $\gamma$ | 0.9 | 0.595 | 0.177 | -0.170 | 0.195 | 0.926 | 0.065 |
| Retrogressive growth $\rho$ | 0.9 | -0.216 | -0.530 | 0.005 | -0.791 | -0.330 | -0.130 |
| Mean recruitment success $\phi$ | 0.9 | -0.381 | 0.453 | 0.093 | 0.763 | -0.626 | 0.056 |
| Degree of iteroparity $S$ | 0.9 | 0.012 | 0.601 | -0.052 | 0.944 | 0.000 | 0.025 |
| Net reproductive rate $R_0$ | 0.9 | -0.053 | 0.255 | 0.529 | 0.609 | -0.052 | 0.757 |
| Mature life expectancy $L_\omega$ | 0.9 | 0.154 | -0.061 | 0.534 | 0.137 | 0.323 | 0.866 |
| Variance | 0.9 | 0.4 | 0.295 | 0.163 | 0.405 | 0.294 | 0.156 |

**Table A2:** PCA loadings at feeding levels of 0.8 and 0.9 for mass corrected non-phylogenetic and phylogenetically informed PCAs. This table show the PC1 values at the feeding level 0.9 as they were prior to inversion.

| Life History Trait | Feeding Level<br>E(Y) | Non-phylogenetic mass corrected PCA |  |  |  | Phylogenetic mass corrected PCA |  |  |  |
| --- | --- | --- | --- | --- | --- | --- | --- | --- | --- |
|  |  | PC1 | PC2 | PC3 | PC4 | PC1 | PC2 | PC3 | PC4 |
| Generation time $T$ | 0.8 | -0.128 | 0.791 | -0.187 | -0.514 | -0.128 | 0.791 | -0.187 | -0.514 |
| Survivorship curve $H$ | 0.8 | 0.088 | 0.268 | 0.033 | -0.932 | 0.088 | 0.268 | 0.033 | -0.932 |
| Age at maturity $L_\alpha$ | 0.8 | 0.046 | 0.133 | -0.968 | 0.177 | 0.046 | 0.133 | -0.968 | 0.177 |
| Progressive growth $\gamma$ | 0.8 | 0.575 | 0.740 | -0.018 | -0.317 | 0.575 | 0.740 | -0.018 | -0.317 |
| Retrogressive growth $\rho$ | 0.8 | -0.954 | 0.050 | 0.068 | -0.160 | -0.954 | 0.050 | 0.068 | -0.160 |
| Mean recruitment success $\varphi$ | 0.8 | 0.481 | -0.852 | -0.135 | 0.037 | 0.481 | -0.852 | -0.135 | 0.037 |
| Degree of iteroparity $S$ | 0.8 | 0.903 | -0.168 | -0.108 | -0.198 | 0.903 | -0.168 | -0.108 | -0.198 |
| Net reproductive rate $R_0$ | 0.8 | 0.342 | -0.311 | -0.873 | 0.006 | 0.342 | -0.311 | -0.873 | 0.006 |
| Mature life expectancy $L_\omega$ | 0.8 | -0.115 | 0.151 | -0.818 | -0.494 | -0.115 | 0.151 | -0.818 | -0.494 |
| Variance | 0.8 | 0.333 | 0.324 | 0.211 | 0.087 | 0.333 | 0.324 | 0.211 | 0.087 |
| Generation time $T$ | 0.9 | 0.423 | 0.195 | 0.800 | 0.289 | 0.423 | 0.195 | 0.800 | 0.289 |
| Survivorship curve $H$ | 0.9 | -0.017 | 0.039 | 0.190 | 0.966 | -0.017 | 0.039 | 0.190 | 0.966 |
| Age at maturity $L_\alpha$ | 0.9 | 0.068 | 0.972 | 0.133 | -0.169 | 0.068 | 0.972 | 0.133 | -0.169 |
| Progressive growth $\gamma$ | 0.9 | -0.307 | -0.060 | 0.915 | 0.131 | -0.307 | -0.060 | 0.915 | 0.131 |
| Retrogressive growth $\rho$ | 0.9 | 0.927 | -0.046 | -0.127 | 0.231 | 0.927 | -0.046 | -0.127 | 0.231 |
| Mean recruitment success $\varphi$ | 0.9 | -0.666 | 0.047 | -0.739 | 0.044 | -0.666 | 0.047 | -0.739 | 0.044 |
| Degree of iteroparity $S$ | 0.9 | -0.919 | 0.018 | -0.147 | 0.191 | -0.919 | 0.018 | -0.147 | 0.191 |
| Net reproductive rate $R_0$ | 0.9 | -0.451 | 0.807 | -0.308 | 0.094 | -0.451 | 0.807 | -0.308 | 0.094 |
| Mature life expectancy $L_\omega$ | 0.9 | 0.115 | 0.816 | 0.115 | 0.518 | 0.115 | 0.816 | 0.115 | 0.518 |
| Variance | 0.9 | 0.362 | 0.284 | 0.188 | 0.122 | 0.362 | 0.284 | 0.188 | 0.122 |

**Table A3:** Kaisers criterion at feeding levels of 0.8 and 0.9 for each of the four PCAs.

|  | **Feeding**<br>**Level** | **PC1** | **PC2** | **PC3** | **PC4** | **PC5** | **PC6** | **PC7** | **PC8** | **PC9** |
| --- | --- | --- | --- | --- | --- | --- | --- | --- | --- | --- |
| Non-phylogenetic PCA | 0.8 | 3.764 | 2.472 | 1.600 | 0.610 | 0.401 | 0.077 | 0.042 | 0.022 | 0.012 |
| Phylogenetic PCA | 0.8 | 14.169 | 6.111 | 2.561 | 0.372 | 0.161 | 0.006 | 0.002 | 0.000 | 0.000 |
| Mass-corrected non-phylogenetic PCA | 0.8 | 2.996 | 2.917 | 1.901 | 0.782 | 0.233 | 0.093 | 0.046 | 0.021 | 0.010 |
| Mass-corrected phylogenetic PCA | 0.8 | 8.979 | 8.511 | 3.615 | 0.611 | 0.054 | 0.009 | 0.002 | 0.000 | 0.000 |
| Non-phylogenetic PCA | 0.9 | 3.603 | 2.658 | 1.464 | 0.827 | 0.283 | 0.106 | 0.040 | 0.010 | 0.008 |
| Phylogenetic PCA | 0.9 | 13.268 | 7.011 | 1.965 | 0.830 | 0.047 | 0.013 | 0.002 | 0.000 | 0.000 |
| Mass-corrected non-phylogenetic PCA | 0.9 | 3.254 | 2.557 | 1.695 | 1.101 | 0.199 | 0.124 | 0.055 | 0.010 | 0.005 |
| Mass-corrected phylogenetic PCA | 0.9 | 10.587 | 6.538 | 2.873 | 1.211 | 0.040 | 0.015 | 0.003 | 0.000 | 0.000 |

**Figure A2:**
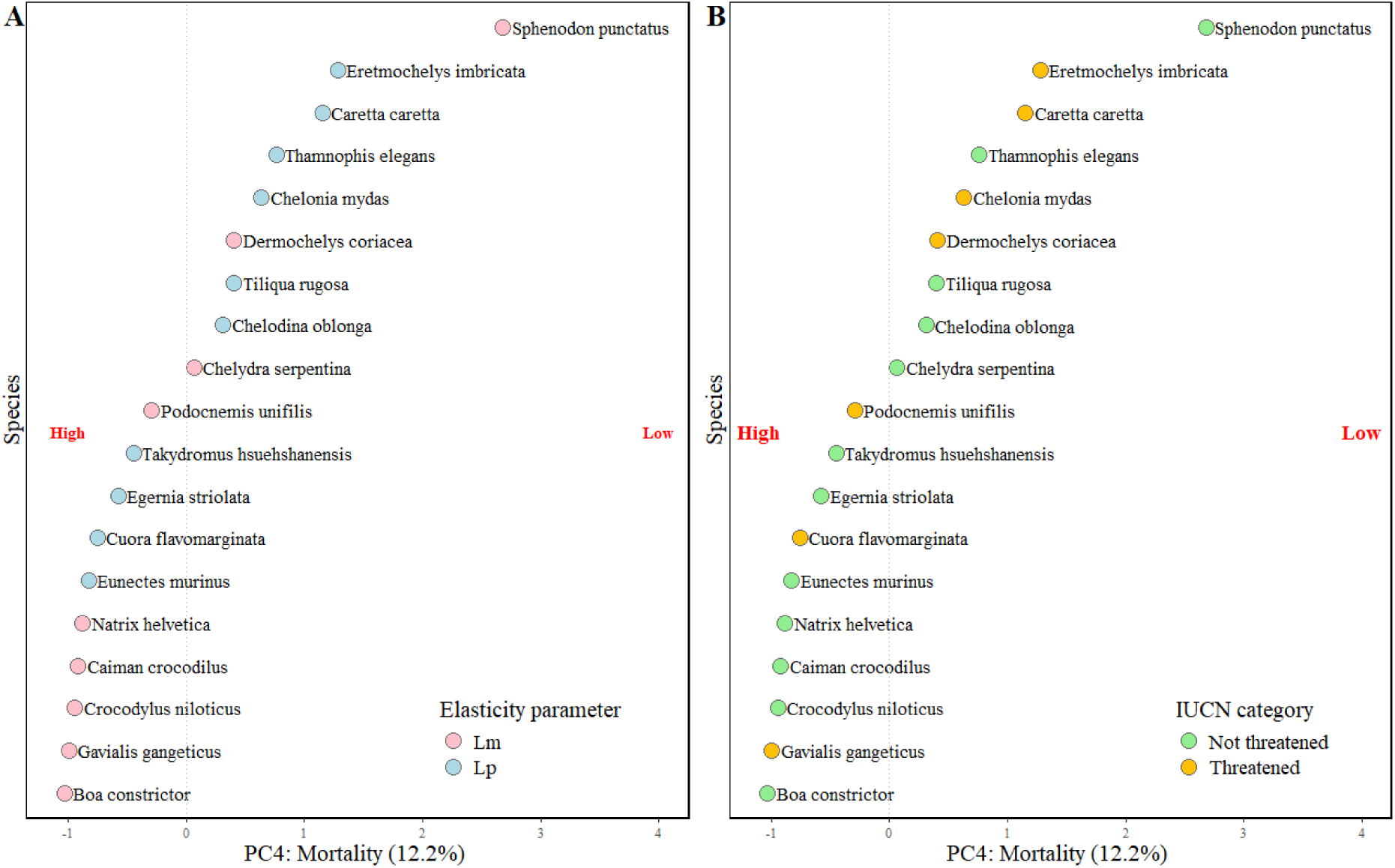
PCA of species position along the most significant life history strategy axis, PC 4 (mortality), coloured by the variable of conservation interest: (A) elasticity and (B) extinction risk. Percentages in brackets shows the percentage variance in life histories explained by the PC axis.

